# Understudied and underprotected: biodiversity and conservation challenges in the Konkan region of Maharashtra

**DOI:** 10.1101/2025.05.16.653652

**Authors:** Kruti Chhaya, Gurudas Nulkar, Parth Bapat, Hrushikesh Barve, Nupur Bhave, Anup Bokkasa, Vaidehi Dandekar, Sachin Desai, Ketaki Ghate, Revati Gindi, Amruta Joglekar, Atul Joshi, Poorva Joshi, Satish Kamat, Manasi Karandikar, Chaitrali Khod, Mandar Kulkarni, Himanshu Lad, Akshay Mandavkar, Pratiksha Mestry, Yash Mishrikotkar, Prateek More, Sayali Nerurkar, Madhura Niphadkar, Ankur Patwardhan, Pooja Pawar, Girish Punjabi, Pratik Purohit, Medhavi Rajwade, Manali Rane, Virashree Raorane, Nikit Surve, Sanjay Thakur, Aparna Watve, Rohit Naniwadekar

**Affiliations:** Nature Conservation Foundation, Mysore; Centre for Sustainable Development, Pune; Konkan Science Forum, Ratnagiri; HRIDAY Energy Network Forum, Pune; The Habitats Trust, New Delhi; Syamantak Trust, Sindhudurg; Oikos for ecological services, Pune; Vanam Ecologics, Pune; Ecovrat Envirosolutions, Pune; FLAME University, Pune; IUCN SSC Western Ghats Plants Specialist Group, Pune; People’s Empowering Movement, Ratnagiri; IUCN Youth Professional Task Force, Pune; Individual Consultant, Pune; Mumbai Tarun Bharat, Mumbai; RANWA, Pune; Sahyadri Sankalp Society, Ratnagiri; Ecological Society, Pune; Wildlife Conservation Trust, Mumbai; Bombay Environmental Action Group, Mumbai; Independent Researcher, Sindhudurg; Wildlife Conservation Society - India, Bengaluru; Adhiwas Foundation, Pune; IUCN Sustainable Use and Livelihoods Specialist Group; IUCN SSC Hornbill Specialist Group

**Keywords:** Infrastructure Development, Forest Loss, Land Use Change, Monoculture Plantations, Western Ghats

## Abstract

In biodiversity hotspots, such as the Western Ghats, periodic synthesis of existing ecological research can help identify knowledge gaps and address critical threats to biodiversity. The Maharashtra part of the Konkan region, situated between the Sahyadri foothills and the west coast, with diverse tropical vegetation, harbours unique open ecosystems, such as lateritic plateaus, and supports a multitude of threatened species, including hornbills and tigers. However, the region remains relatively understudied and has not received adequate protection through state or national conservation policies and laws. The area is undergoing rapid human-driven land-use change. These activities can impact the region’s biodiversity, necessitating an effort to identify knowledge gaps and critical threats. Through a combination of a literature review and focus group discussions with 44 participants from various institutions and non-governmental organisations, we synthesised existing published information on the Konkan region of Maharashtra and identified key research gaps and conservation challenges. Our review of 138 studies found that while agroforestry and human-wildlife interactions have received some research attention, the effects of climate change on the region’s biodiversity remain poorly understood. Focus group discussions highlighted major threats, including land-use changes due to expanding monoculture plantations, clear-felling of forests, forest fires, environmental pollution, and rapid infrastructure development, which are leading to habitat loss and fragmentation. Some of these concerns were validated by publicly available regional data, which revealed a 30% increase in roads, a 14% expansion of cashew plantations, and associated forest loss. This synthesis offers valuable insights for government and non-governmental organisations to inform future research and conservation efforts in the region.

## 1 INTRODUCTION

The Western Ghats-Sri Lanka Biodiversity Hotspot is recognised as one of the world’s highest conservation priority biodiversity hotspots due to its high endemism and the threats faced by its biodiversity and ecosystems (Myers et al 2000; Mittermeier et al 2011). More than 33,500 km^2^ of forest was lost between 1920 and 2013 in the Western Ghats region of the hotspot (Reddy et al 2016). Expanding coffee, tea, rubber, cashew, and mango plantations, along with other forms of agriculture and forest degradation, are the primary drivers of this forest loss (Reddy et al 2016). While significant scientific information exists from the central and southern Western Ghats, the northern Western Ghats have received relatively less research and conservation attention despite being geologically and biogeographically distinct from the central and southern Western Ghats and facing high threat levels (Srivathsa et al 2023).

The northern Western Ghats lie between 14° (north of Kali River, Karnataka) and 21°N (south of the Tapi River, Gujarat) (Subramanyam and Nayar 1974). Their unique geography becomes apparent as one moves west towards the coast from the Sahyadri mountains. This unique geography is a consequence of Deccan volcanic activity around 60 million years ago, followed by weathering, which resulted in distinctive lateritic plateaus and relatively steep escarpments of the Sahyadris (Figure 1). Within the northern Western Ghats, the Konkan region of Maharashtra is a distinct physiographic region, spanning a width of 40 to 50 km between the coastline to the west and the escarpment of the Western Ghats to the east.

**Figure 1.**
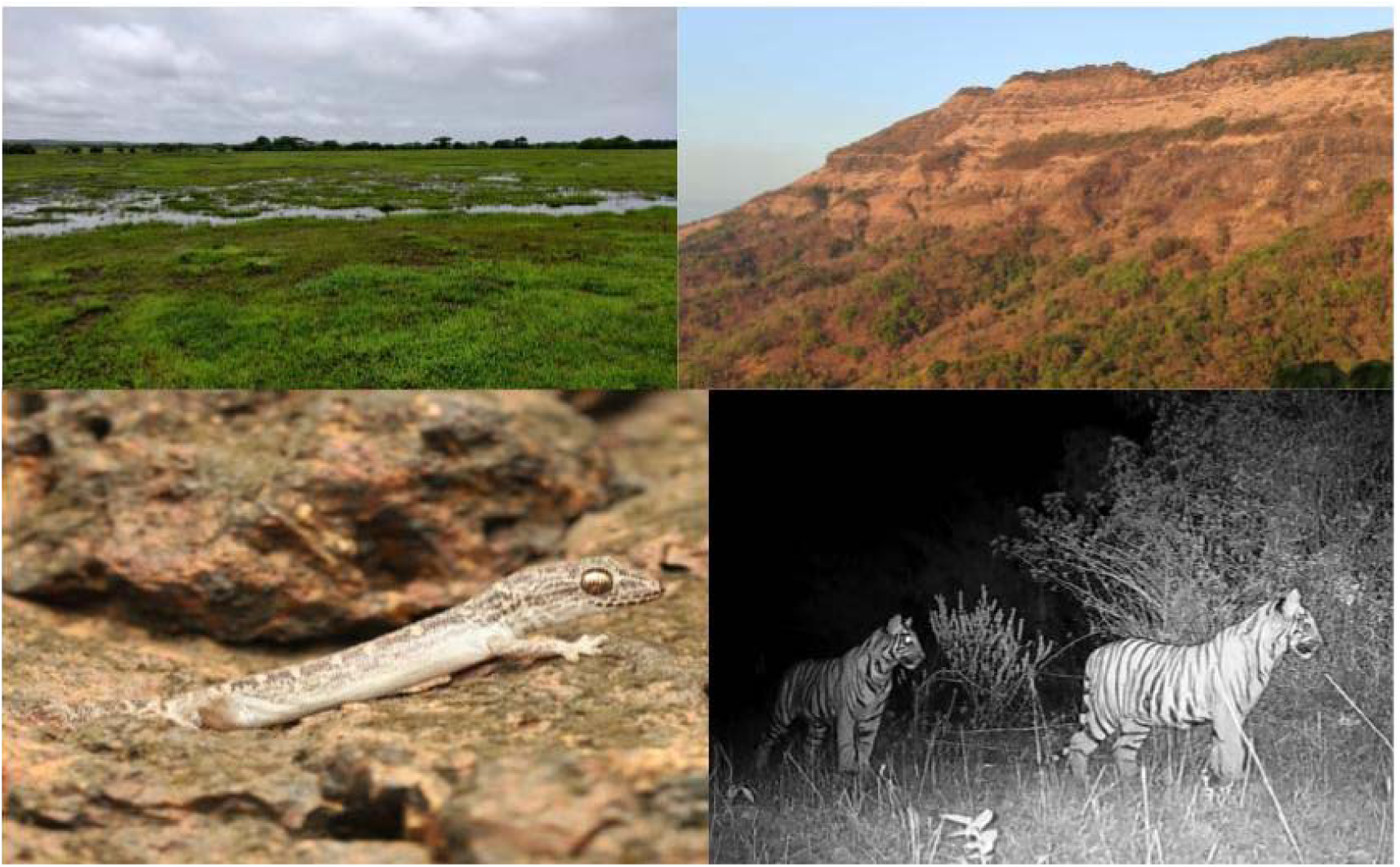
Lateritic plateau (Top left) and steep escarpments of the Sahyadris (Right). White-striped Viper Gecko, *Hemidactylus albofasciatus*, found only at select plateaus in Ratnagiri (Bottom left) and tiger cubs image captured through camera traps (Bottom right). Photos: Aparna Watve, Rohit Naniwadekar, WCT/Maharashtra Forest Department.

Large lateritic plateaus, deeply entrenched river channels, and piedmont plains at the foot of the Sahyadri escarpment are significant land features, particularly in the Ratnagiri and Sindhudurg Districts (Diddee et al 2002). The past geological events have significantly shaped the biogeography of the region (Joshi and Karanth 2013). Unlike forested ecosystems, the lateritic plateaus are exposed to the sun for nearly five to six months a year and are lashed by heavy monsoon rainfall. This extreme climate has fostered a diverse array of herbaceous plants and herpetofauna adapted to these harsh environments. A significant proportion of this diversity is endemic to the region. Seventeen percent of the total herbaceous plant species on low-level lateritic plateaus are endemic to the region (Kulkarni et al 2023). *Dipcadi concanensis* and *Iphigenia ratnagirica*, classified as Endangered by IUCN Red List have large populations on the lateritic plateaus in Ratnagiri and Sindhudurg. Species such as the White-striped Viper Gecko (*Hemidactylus albofasciatus)* (Figure 1), classified as Vulnerable by the IUCN, are restricted to select lateritic plateaus in the Ratnagiri District (Amberkar and Mungikar 2024; Jithin et al 2023). The low-elevation forests of the Konkan region harbour significant populations of threatened fauna, including hornbills (Biswas et al 2023; Pawar and Sadekar 2023), tigers, gaur, elephants, and other wildlife (Punjabi and Rao 2017; Rege et al 2020) (Figure 1). Moreover, the geographic range of several endemic species of the Western Ghats, for example, the Western Ghats King Cobra (*Ophiophagus kaalinga*), Draco (*Draco dussumieri*), Hump-nosed Pit Viper (*Hypnale hypnale*), Malabar Woodshrike (*Tephrodornis sylvicola*), Grey-headed Bulbul (*Microtarsus priocephalus*), extend into the Konkan region of Maharashtra, underscoring its biogeographic relevance.

While agroforestry plantations are prevalent across the northern, central, and southern Western Ghats, numerous studies have compared biodiversity responses across agroforestry plantations and forests in the central and southern Western Ghats (Anand et al 2008; Daniels et al 1990; Karanth et al 2016; Raman and Sukumar 2002; Raman et al 2021; Ranganathan et al 2010; Sidhu et al 2010), yet there are very few peer-reviewed publications from the Konkan region of the northern Western Ghats (Jithin et al 2023; Jithin and Naniwadekar 2024; Lad et al 2024). There are very few Protected Areas between 18.5°N and 15.6°N in the Konkan region of Maharashtra; the majority of the Protected Areas within these latitudes (Sahyadri Tiger Reserve, Radhanagari and Tamhini Wildlife Sanctuaries, and a series of Conservation Reserves) are mostly restricted to the crest of the Western Ghats (locally called “*Desh*”). Among other land use changes, the lateritic plateaus and foothill forests under private ownership in Konkan are being converted into monoculture plantations of mango, cashew, and rubber at an alarming rate. Impacts of these plantations on biodiversity are understudied. Despite the Konkan region’s unique biogeography and biodiversity value and the threats it faces, it has not been identified as a critical area of conservation importance (Srivathsa et al. 2023), a likely consequence of relatively poor representation of the literature from the area.

Given this background, we deemed it critical to assess the existing knowledge of anthropogenic impacts on biodiversity and identify critical knowledge gaps to prioritise future research and conservation efforts in the Konkan region. Our objectives were to 1) review the existing literature on biodiversity and conservation in the Konkan region, 2) identify key anthropogenic threats to biodiversity, and 3) assess temporal change in select anthropogenic threats at a regional scale using data available in the public domain. While the first objective encompassed the entire Konkan region in Maharashtra, objectives 2 and 3 focused exclusively on the Ratnagiri and Sindhudurg Districts of Maharashtra, as these areas continue to harbour significant natural habitat and biodiversity but lack extensive Protected Area coverage (Figure 2). Moreover, the districts around Mumbai (Thane, Palghar, and Raigad) have experienced land-use change and industrialisation for more than four decades, while Ratnagiri and Sindhudurg have experienced rapid land-use change in the last two decades. While the first objective relied on synthesising insights from published literature, the second objective sought information from experts from the region and their field experiences to identify key threats, and the third objective involved analysing publicly available information related to certain threats.

**Figure 2.**
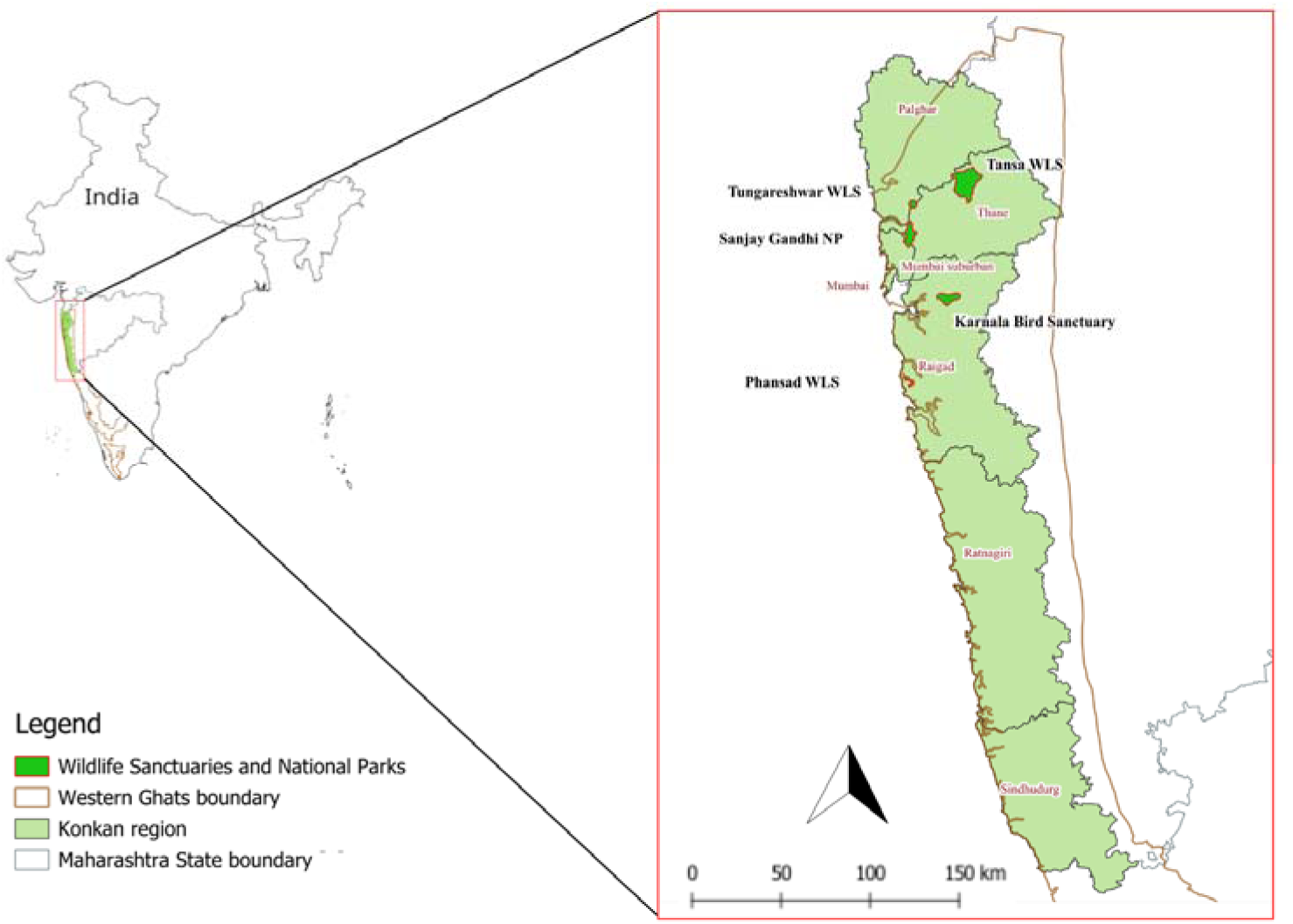
Indicative boundaries of the Wildlife Sanctuaries and National Parks in the Konkan region of Maharashtra. (India Biodiversity Portal, Survey of India website). NP: National Park, WLS: Wildlife Sanctuary.

## 2 METHODS

### 2.1 Study Area

The Konkan (or Kokan) region of Maharashtra, India, is a 40-50 km wide coastal strip bounded by the northern Western Ghats to the east and the Arabian Sea to the west. Covering an area of 30,728 km^2^, it has a population of 28.6 million as per the 2011 Census. Administratively, the region includes the districts of Mumbai, Thane, Palghar, Raigad, Ratnagiri, and Sindhudurg (Figure 2). Physiographically, Konkan is divided into north, middle, and south sections, each distinguished by unique physical, geological, environmental, and cultural features. The Ratnagiri and Sindhudurg Districts form the southern Konkan. The region experiences a seasonal climate. Summers (March-early June) are hot and humid, with temperatures reaching 33, while winter (November-February) are relatively mild, with minimum temperatures around 18. The annual average rainfall in the region is 3,225 mm, primarily due to the south-west monsoon from June to September. The 2012 Maharashtra Human Development Report (Planning Department 2012) ranked Sindhudurg as ‘very high’ and Ratnagiri as ‘high’ on the Human Development Index (HDI), with both districts performing well in education and health indicators. Additionally, th 2022 Directorate of Economics and Statistics (DES) report (DES 2022) highlights good road connectivity to the villages across Sindhudurg and Ratnagiri.

### 2.2 Literature Review

We conducted a literature review focusing on the districts of Palghar, Thane, Mumbai, Raigad, Ratnagiri, and Sindhudurg in the Konkan region of Maharashtra. Using a combination of taxa, threat, region, and habitat-related keywords, we searched Google Scholar for ecological and conservation science literature on plants and vertebrates from terrestrial ecosystems. We systematically scanned Google Scholar results pages until no further relevant literature was found. The keywords used for the review are listed in Table S1. Since we focused on ecological or conservation science studies, we excluded literature limited to species checklists or natural history observations. We collated all the literature available that we could find on Google Scholar (since we did not have access to Web of Science) till 7^th^ July 2024, with the oldest publication dating back to 1981.

The collected literature was classified into peer-reviewed publications, reports, preprints, theses, book chapters, and conference papers. We recorded the district(s) where each study was conducted and noted whether the author’s affiliation was a non-governmental organisation (not-for-profit organisation), a college or university, or a research institute. We classified Research institutes separately, as, unlike colleges and universities, the primary mandate of a research institute is research, not teaching. The focal taxa were categorised as plants, amphibians, reptiles, birds, and/or mammals, and we identified whether studies were conducted inside and/or outside Protected Areas. Additionally, we classified the literature based on relevant threat categories that emerged from the expert interactions and Focus Group Discussions.

### 2.3 Focus group discussions for identifying threats to biodiversity in Ratnagiri and Sindhudurg

To complement the findings from our systematic literature review, we conducted a day-long, in-person meeting with key informants at the Gokhale Institute of Politics and Economics, Pune, India, on 13^th^ July 2024. This meeting aimed to review studied threats and identify threats that may not have been studied in existing published literature. We invited academics, conservationists, activists, journalists, and lawyers involved in nature conservation in the Konkan region. Given their involvement in research and conservation within the area, they were well-positioned to provide broader insights into the nature of existing threats. The focus of the discussion was on the Ratnagiri and Sindhudurg Districts, which, from a biodiversity perspective, still contain large forest and natural lateritic plateau habitats that are unprotected and outside the network of Protected Areas in the Konkan region.

During the meeting, participants were divided into four groups to list the threats to terrestrial biodiversity in the Ratnagiri and Sindhudurg Districts of the Konkan region. Each group then discussed whether these threats 1) have increased or decreased over the past 20 years, 2) are localised to specific sites or widespread across Ratnagiri and Sindhudurg Districts, and 3) whether there is existing knowledge of their impacts on biodiversity in the region. The findings of each group were synthesised, and threats were prioritised based on their geographic extent, perceived trends over time, and the availability of existing knowledge on their impacts. For instance, widespread threats, perceived as increasing, and having limited background knowledge were ranked as having higher priority for future research and conservation action. For some threats, we also triangulated participants’ perceptions with existing data from different sources, as outlined in the next section.

### 2.4 Determining temporal change in select threat categories

We determined temporal changes between 2009 and 2023 in threat categories that emerged as priorities in the Focus Group Discussions and for which temporal data were available publicly. We extracted information from published data available on government websites and peer-reviewed publications on the following themes: 1) road length, 2) number of mines (bauxite, iron ore), 3) extent of land under specific agroforestry plantations (e.g., cashew, mango), and 4) primary forest loss in the Ratnagiri and Sindhudurg Districts.

We obtained data on the total road length (including national and state highways, district roads, and village roads) in the Ratnagiri and Sindhudurg districts, as well as the number of bauxite mines in Ratnagiri from the Infrastructure Statistics of Maharashtra State reports, published by the Directorate of Economics and Statistics (DES 2009-2023). However, we found no information on bauxite mines for Sindhudurg. Similarly, we retrieved data on the number of iron ore mines in Sindhudurg, but no corresponding information was available for Ratnagiri. Data on the area under mango and cashew cultivation in Ratnagiri and Sindhudurg were sourced from District Social and Economic Reviews (DES 2010–2023a, 2010-2023b) and HAPIS Area Production Estimate Reports (Ministry of Agriculture and Farmers Welfare (MoAFW) 2013– 2023). Additionally, we used data from Global Forest Watch (University of Maryland and World Resources Institute 2024) to estimate primary forest loss in these districts from 2009 to 2023. The details of data sources for the threats mentioned above are provided in Table S2. We then calculated the percentage change in these activities for each region over time.

## 3 RESULTS

### 3.1 Literature Review

We identified 138 publications authored by 95 unique first authors affiliated with 61 unique organisations (Table A3). While 39% of these studies were conducted in a single district, 34% spanned multiple districts, and the remaining studies were not site-specific. At least 35.5% of the studies were conducted entirely or partly in Sindhudurg District, 31.8% in Ratnagiri, 17.3% in Raigad, 13.7% in Thane, 17.3% in Mumbai and 5% in Palghar. The first authors were primarily affiliated with colleges or universities (40%), non-governmental organisations (38%), and research institutes (18%). The number of publications has increased over time (Figure 3). Of the 138 publications, 71% (98) were peer-reviewed scientific articles published in 58 different scientific journals, 13% (18) were reports, 6% (8) were preprints, and 4% (5) were doctoral theses. The rest were master’s theses, book chapters and conference papers. Most of the studies focused on plants (35%), followed by mammals (22%), birds (17%), amphibians (12%), and reptiles (7%). Over 55% of the studies were conducted in non-protected areas, approximately 14% in protected areas, and the remainder in both protected and non-protected areas.

**Figure 3.**
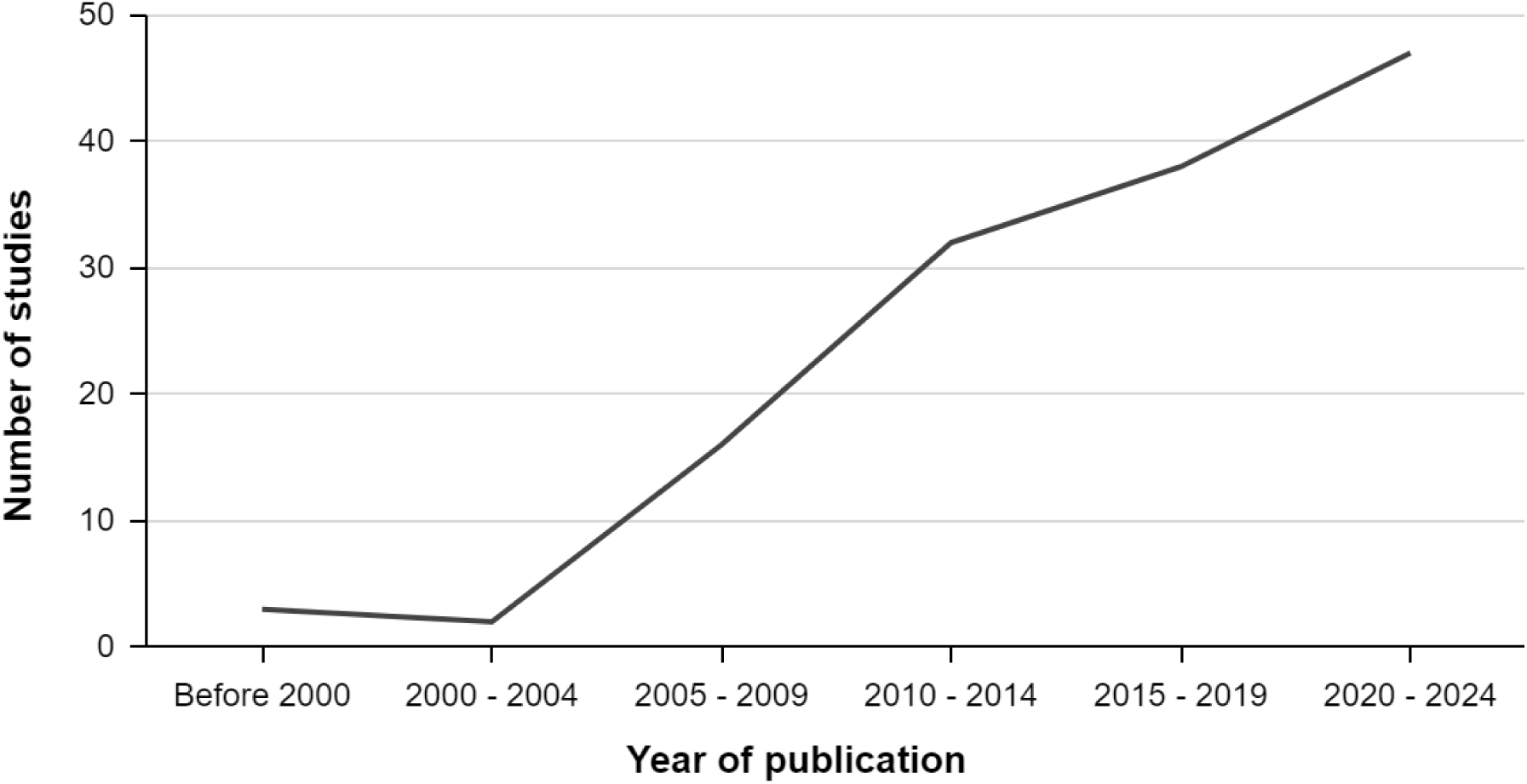
Based on Google Scholar search results, the number of ecological and conservation science studies from the Konkan region published over time. Over the last two decades, a consistent increase in the number of studies has been seen. Table S3 includes details of all studies represented in the figure.

Of the 138 studies, we classified 85 studies as ‘conservation science’ studies, which evaluated the effects of human activities on biodiversity, and 53 as ‘ecological’ studies, which primarily aimed to investigate the ecology of wildlife species or communities. We categorised the conservation science studies into 24 categories (Table 1). The most common topics were the impacts of agroforestry (n = 14) and human-wildlife interactions (n = 14), while the effects of climate change were addressed by only one study. Table 1 summarises the number of studies investigating the impacts of various anthropogenic activities across these categories.

**Table 1.** The number of studies investigating the impacts of different human activities on biodiversity. Please refer to Table S3 for relevant studies categorised under different sections.

| Sr. No. | Human Activity | Number of Studies |
| --- | --- | --- |
| 1 | Agroforestry (Mango, Cashew, Rubber) | 14 |
| 2 | Human-wildlife interaction | 14 |
| 3 | Mining and quarrying | 7 |
| 4 | Logging and commercial harvest of plant resources (fuelwood collection) | 6 |
| 5 | Livestock: grazing, trampling | 5 |
| 6 | Tourism | 4 |
| 7 | Urbanisation | 4 |
| 8 | Agriculture | 3 |
| 9 | Fire | 3 |
| 10 | Fragmentation and forest loss | 3 |
| 11 | Transportation/ roads | 3 |
| 12 | Hunting | 3 |
| 13 | Infrastructure development (refinery, ports) | 2 |
| 14 | Pollution: industrial effluents, air pollution | 2 |
| 15 | Wildlife trade | 2 |
| 16 | Blasting | 1 |
| 17 | Breakdown of traditional management systems | 1 |
| 18 | Climate change | 1 |
| 19 | Construction | 1 |
| 20 | Cultural activities | 1 |
| 21 | Free-ranging dogs | 1 |
| 22 | Invasives | 1 |
| 23 | Windmill | 1 |
| 24 | Starvation of wildlife due to anthropogenic changes | 1 |

### 3.2 Focus group discussion outputs

Forty-four participants from 27 institutes attended the stakeholder meeting, including two academicians, 38 conservation scientists and/or practitioners, one media representative, two environmental law professionals, and one corporate sustainability professional, were divided among four groups. The main threats identified by the four groups are listed in Table 2. We categorised these threats into those that primarily cause drastic changes in land use, pollute the environment, or impact biodiversity and ecosystem function (Table 2). Discussions highlighted human activities with limited information on their biodiversity impacts (e.g., illegal wildlife trade, harvest of medicinal plants or wild vegetables) (Table 2). They also revealed a lack of regional-level information on several potential threats, some of which may be critical for determining sustainability thresholds. Many potential threats are widespread in the region and appear to be increasing over time.

**Table 2.** Summary of threats to biodiversity in the Ratnagiri and Sindhudurg Districts of Maharashtra as identified by 44 key informants during the in-person meeting at Gokhale Institute of Politics and Economics, Pune, on 13^th^ July 2024. The table categorises each threat based on its geographic scale (local or regional), temporal trend (increasing, decreasing, or stable), and whether its impacts on biodiversity have been studied in the region. For ease of organisation, the activities have been grouped into six categories as shown in the extreme right column.

| Sr.<br>No. | Potential Threats | Local/<br>Regional | Increasing/<br>Decreasing/ Stable | Existing<br>Information of<br>Impacts on<br>Biodiversity | Category |
| --- | --- | --- | --- | --- | --- |

| **Sr. No.** | **Potential Threats** | **Local/ Regional** | **Increasing/ Decreasing/ Stable** | **Existing Information of Impacts on Biodiversity** | **Category** |
| --- | --- | --- | --- | --- | --- |
| 1 | Road widening, highways | Regional | Increasing | No | Land use |
| 2 | Monoculture plantations (e.g., rubber, cashew, mango) | Regional | Increasing | Some | Land use |
| 3 | Clearfelling of private forests for fuelwood | Regional | Increasing | Some | Land use |
| 4 | Forest diversion for infrastructure projects (e.g., hydel projects, mines, etc.) | Regional | Increasing | Some | Land use |
| 5 | Commercial plotting | Regional | Increasing | Some | Land use |
| 6 | Urbanisation | Regional | Increasing | Some | Land use |
| 7 | Degradation and loss of sacred groves | Regional | Increasing | Yes | Land use |
| 8 | Mining/quarrying | Local | Increasing | No | Land use |
| 9 | Infrastructure development (e.g., refinery, ports) | Local | Increasing | No | Land use |
| 10 | Agricultural pollution (e.g., pesticides, weedicides) | Regional | Increasing | Some | Pollution |
| 11 | Industrial pollution | Regional | Increasing | Some | Pollution |
| 12 | Solid waste management, especially in rural areas (e.g., plastic) | Regional | Increasing | Some | Pollution |
| 13 | Unregulated tourism | Regional | Increasing | Yes | Pollution |
| 14 | Fire incidents | Regional | Increasing | No | Impacts on biodiversity and ecosystem |
| 15 | Extreme climate events | Local and Regional | Increasing | Yes | Impacts on biodiversity and ecosystem |
| 16 | Human-wildlife interactions | Regional | Increasing | Some | Impacts on biodiversity and ecosystem |
| 17 | Habitat degradation | Regional | Increasing | Some | Impacts on biodiversity and ecosystem |
| 18 | Invasive plants | Regional | Increasing | Yes | Impacts on biodiversity and ecosystem |
| 19 | Bushmeat hunting | Regional | Decreasing | No | Impacts on biodiversity and ecosystem |
| 20 | Illegal wildlife trade | Do not know | Do not know | No | Impacts on biodiversity and ecosystem |
| 21 | Harvest (e.g., wild vegetables, medicinal plants) | Do not know | Do not know | No | Impacts on biodiversity and ecosystem |
| 22 | Livestock grazing | Local | Do not know | No | Impacts on biodiversity and ecosystem |
| 23 | Planning and management given to state bodies while excluding local bodies | Regional | Increasing | No | Policy and property |
| 24 | Poor management of sensitive areas | Local and Regional | Increasing | Yes | Policy and property |
| 25 | Misinterpretation by policymakers | Local and Regional | Mixed | No | Policy and property/Social issues |
| 26 | Complicated or lack of clarity in property ownership | Regional | Stable | Some | Policy and property |
| 27 | Lack of demarcation of Common Property Resources (e.g. water bodies - high flood line and buffer | Regional | Stable | Some | Policy and property |

| Sr. No. | Potential Threats | Local/ Regional | Increasing/ Decreasing/ Stable | Existing Information of Impacts on Biodiversity | Category |
| --- | --- | --- | --- | --- | --- |
|  | zone) |  |  |  |  |
| 28 | Policy (e.g., Private Forest Acquisition Act) | Regional | Stable | Some | Policy and property |
| 29 | Breakdown of traditional management systems | Do not know | Do not know | Do not know | Policy and property |
| 30 | Lack of environmental awareness, especially among stakeholders (e.g., politicians and citizens) | Regional | Increasing/Mixed | No | Community engagement and awareness |
| 31 | Scientific findings not communicated to locals | Local and Regional | Mixed | No | Community engagement and awareness |
| 32 | Spread of misinformation (e.g., fake news) | Local and Regional | Mixed | No | Community engagement and awareness |

### 3.3 Determining rates of increase in certain human activities

By collating information from government records and other sources on regional (e.g., road expansion, agroforestry plantations) and local (e.g., mining) threats, we found that the road network in Ratnagiri and Sindhudurg Districts has increased by more than 30% since 2009 (Table 3). Since 2013, the area under cashew cultivation has grown by more than 14% in the two districts (Table 3). At the site level, the number of bauxite mines in Ratnagiri District rose from three to eight since 2009, while iron ore mines in Sindhudurg increased from three to nine between 2009 and 2023. Concurrently, up to 2.3% of primary forests were lost during a similar period since 2009 (Table 3). A district-wise summary for each of the threats listed above is presented in Table 3.

**Table 3.**
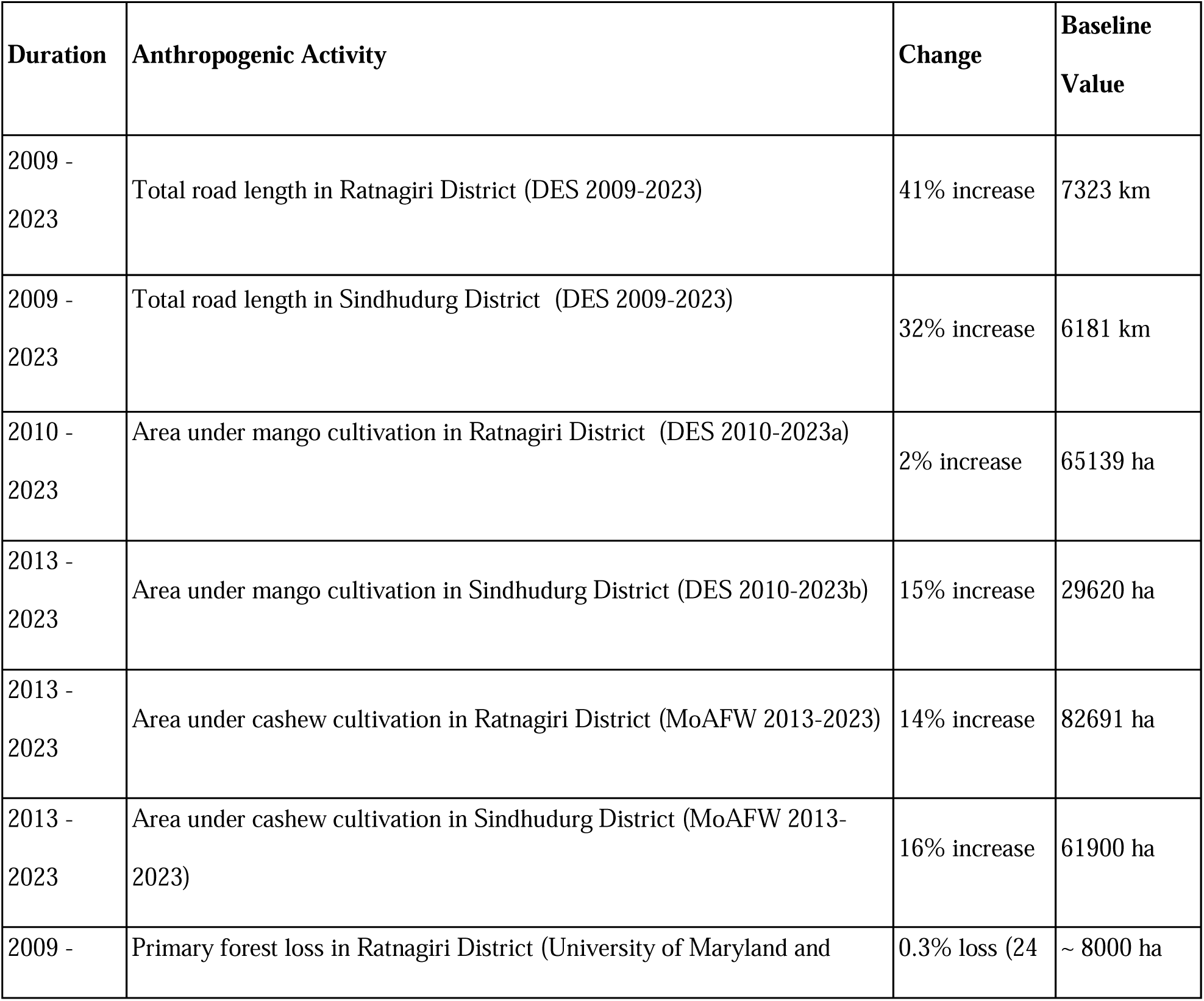
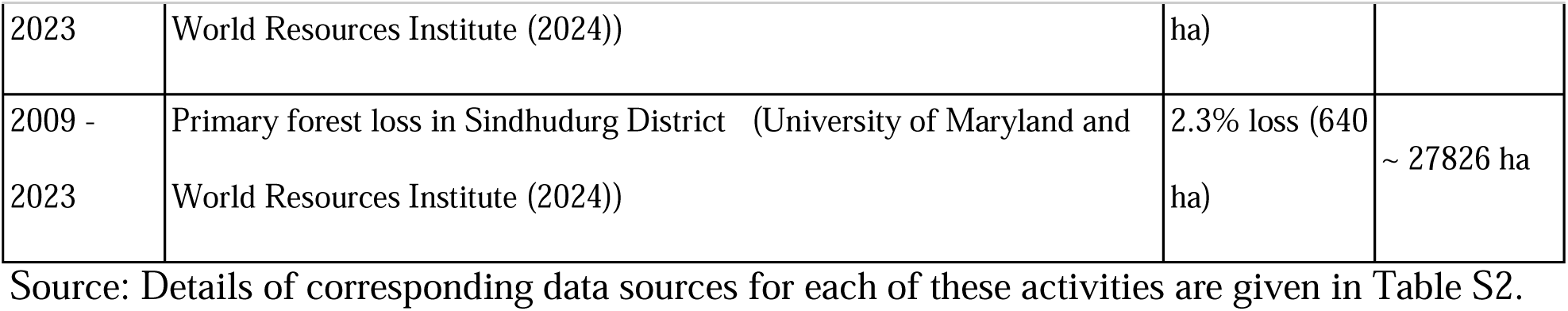
Summary of percent change in two regional-level human activities (e.g., roads, agroforestry) over time. The table includes the periods for which changes were assessed and the baseline value for each activity.

## 4 DISCUSSION

In this study, we integrated information from three approaches—literature review, focus group discussions, and data gleaned from public sources—to identify and prioritise human activities threatening biodiversity in the Konkan region of Maharashtra. While the number of ecological and conservation science studies is steadily increasing, they do not fully address the research and action needs. These identified knowledge gaps must be addressed in the near future to effectively mitigate the impacts of these threats on biodiversity and ecosystem function. We found that herpetofauna, which exhibits high diversity and endemism in the Western Ghats, are relatively under-represented in the published literature. Interestingly, not-for-profit organisations play a pivotal role in knowledge generation. Published information is mostly available as scientific papers and preprints. In contrast, reports are relatively less accessible, underscoring the need to publish them on open-source platforms (e.g., Zenodo). The group discussions revealed more than 30 regional and local-level factors across various themes, including land-use change, pollution, policy, and community engagement and awareness issues. Unfortunately, actionable information for many of these factors remains lacking. Consistent with observations from focus group discussion participants, secondary sources provided evidence of increasing regional and local-level threats to biodiversity in the Ratnagiri and Sindhudurg districts, such as expanding road networks, agroforestry plantations, and mining activities. Additionally, secondary sources revealed evidence of forest cover reduction, particularly in the Sindhudurg district of Maharashtra.

### 4.1 Identifying knowledge gaps

While the number of studies assessing the impacts of various human activities on biodiversity is increasing over time, our focus group discussions revealed pervasive threats for which little regional information is available. These include activities such as infrastructure development, fires, illegal wildlife trade, and bushmeat hunting, among others. These activities can significantly impact biodiversity and ecosystem function (Bashyal et al 2021; Evans et al 2020; Feng et al 2021; Spencer et al 2023). However, detailed regional-level data on the intensity of these impacts and their geographic extent are still needed to inform future conservation planning in the region.

The impacts of agroforestry on biodiversity, which were identified to be rapidly spreading in the landscape, are relatively well studied in the region. There is information on the responses of multiple taxa, including mammals (Rege et al 2020), birds (Biswas et al 2023; Madhu et al 2024; Munje and Kumar 2022), and amphibians (Gosavi et al 2025; Lad et al 2024). However, the role of remnant forest patches and native trees in agroforestry plantations in influencing biodiversity and critical ecosystem functions, such as pollination, which may benefit agroforestry plantations, needs to be better understood. This information will be essential for shaping agroforestry and biodiversity conservation policies in this human-dominated landscape. It is essential to explore how the ongoing expansion of monoculture plantations can be made more biodiversity-friendly through land-sharing or land-sparing approaches (Birch et al 2024; Finch et al 2019). Additionally, since plantations benefit from adjacent forest patches, through enhanced pollination and improved yield (Freitas et al 2014), there is a pressing need for ecological restoration of degraded government- and privately-owned forests. Given the scarcity of primary forest remnants that can serve as a source of seeds and the historical disturbances that have altered forest structure and plant composition (Biswas et al 2024; Kulkarni et al 2013), active restoration efforts may be necessary to support biodiversity recovery.

Studies focusing on the impacts of climate change on biodiversity and ecosystem function for the Konkan region were extremely underrepresented in the literature. The northern Western Ghats is biogeographically distinct from the central and southern Western Ghats due to climatic instability in the past and greater present-day climatic seasonality (Joshi and Karanth 2013). The region features unique habitats, such as lateritic plateaus, where the high diversity and endemism of herbaceous plants (Kulkarni et al 2023; Watve 2013) and other unique biodiversity are strongly linked to climate (Jithin and Naniwadekar 2024). Similarly, the evergreen forests of the northern Western Ghats experience greater rainfall seasonality compared to those in the southern Western Ghats, rendering them more vulnerable to climate variability. The region is experiencing extreme climate events (e.g., excessive rainfall) and is predicted to be highly vulnerable to climate change (Chaturvedi et al 2011). Therefore, long-term studies documenting the impacts of climate change on biodiversity and ecosystem function are critical.

Climate change has been documented to impact traditional livelihoods in the region (ACWADAM 2022). Similarly, the livelihoods of people are also affected by negative human-wildlife interactions, such as crop depredation and loss of human life, which likely result in negative perceptions towards wildlife in the region, an aspect that requires greater attention. This can have cascading impacts on biodiversity persistence in the region. While biodiversity can impact socio-ecological outcomes for people, the reverse is also true. There is also a need for socio-ecological studies that examine the links between market forces (e.g., large-scale fuelwood harvest through clear-felling) and habitat degradation or modification, along with their cascading effects on biodiversity. Understanding these connections could facilitate the development of more permanent and effective solutions to biodiversity threats. Similarly, the drivers behind societal transformations and the consequent breakdown of biodiversity-friendly traditional systems (e.g., degradation of sacred groves) need to be determined to explore possibilities for reviving such systems. The need for additional socio-ecological studies has been identified as a knowledge gap through the focus group discussions and the literature review.

### 4.2 Some up-and-coming threats

#### Linear intrusions

The focal group discussions identified linear intrusions, particularly highways, as a significant threat to biodiversity in the region. Publicly available data also confirmed this, showing more than 30% increase in the road network in the region over the past decade. While there is significant information on the impacts of roads and highways on biodiversity globally, our literature review revealed a lack of such data specific to the Konkan region. This underscores the need for additional studies on the effects of existing roads in the region on habitat fragmentation, wildlife mortality, and animal movement in the area, especially since the area harbours threatened species like Gaur (*Bos gaurus*), Sambar (*Rusa unicolor*), Wild Dog (*Cuon alpinus*), Leopard (*Panthera pardus*) and Tiger (*Panthera tigris*). Such information will be crucial for guiding the future development of linear intrusions in the region. Linear intrusions, particularly roads, are known to increase habitat fragmentation, edge effects, and wildlife mortality, while impeding animal movement, with effects pervading up to 17 km from major roads (Jayadevan et al 2020; Nayak et al 2020) (Figure 4). In the Konkan region, linear intrusions are already causing either channelled or restricted movement of large mammals (Jayadevan et al 2020; Nayak et al 2020). Multiple forest areas in the Konkan region have been categorised as large, unfragmented patches (Jayadevan et al 2020; Nayak et al 2020), which must be protected from further fragmentation due to the expanding road network, given their importance for biodiversity, local communities, and function (e.g., as a source of freshwater, natural resources, preventing natural disasters like landslides). Studies recommend avoiding further fragmentation of forest patches in the Western Ghats by aligning linear infrastructure along existing features and prioritising the north-south axis over the east-west axis (Nayak et al 2020). However, proposed plans to widen existing highways, such as the six-lane Shaktipeeth highway connecting Nagpur and Goa (ET online 2025) or the Konkan Greenfield expressways, will likely exacerbate wildlife fragmentation.

**Figure 4.**
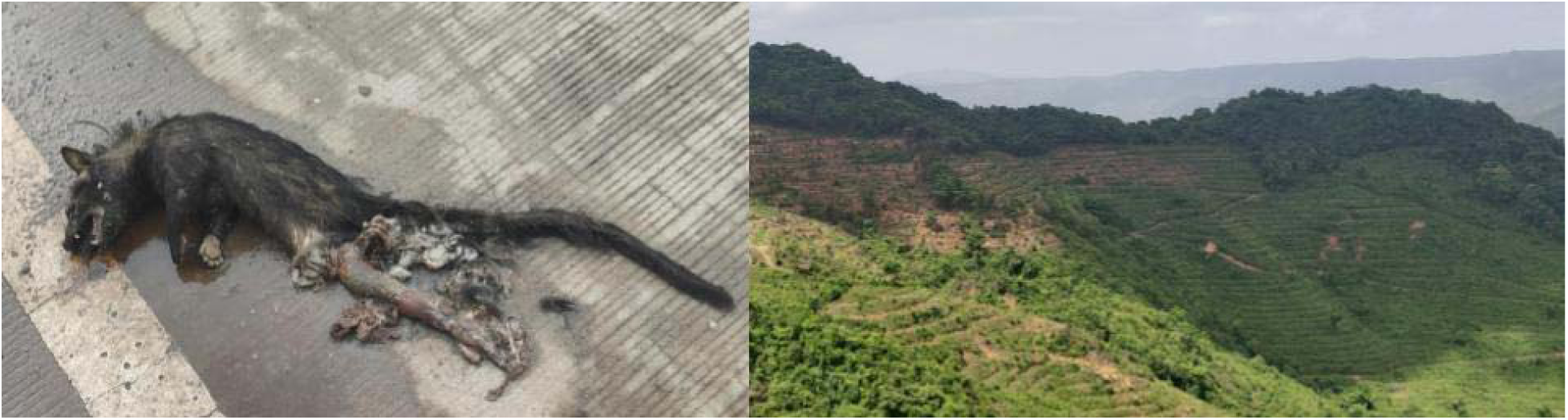
Asian Palm Civet, *Paradoxurus hermaphroditus*, roadkill on Mumbai-Goa highway near Zarap, Sindhudurg (Left) and privately-owned forests cleared, terraced and converted to cashew plantations at Terwan-Medhe in Sindhudurg (Right). (Photo: Rohit Naniwadekar).

#### Human-wildlife interactions

Another challenge identified during the focus group discussions was human-wildlife interactions. While human-elephant interactions in Sindhudurg are documented in the literature (Mehta and Kulkarni 2013; Patil and Patil 2019) no published studies were found on human-gaur interactions in the Konkan region, although some information exists from near Radhanagari Wildlife Sanctuary (Joshi 2008). Given the extensive cover of mango orchards, coconuts and areca nut in the region, primates and squirrels, especially the Indian Giant Squirrel (*Ratufa indica*), often come in conflict with humans due to crop damage. However, the intensity and extent of its impact on landowners are poorly understood. Similarly, impacts of these orchards on primates and squirrels are poorly understood, although an unpublished study indicates reduced densities of langurs and squirrels in monoculture plantations compared to forests. Elephant movement and range extension from Karnataka to Maharashtra and Goa was first recorded in 2002. Studies have reported an increase in crop raiding incidents by elephants in Sindhudurg from 2002 to 2015 (Mehta and Kulkarni 2013; Patil and Patil 2019). In 2015, 33% of the villages in Sindhudurg were affected by crop raiding by elephants (Mehta and Kulkarni 2013). Given the expansion of agroforests and the reduction in forest cover in the region, these interactions are likely to intensify. There is a need for enhanced community engagement and capacity building for Forest Department staff to manage the increasing interactions between humans and large mammals. This would support biodiversity persistence in these human-dominated landscapes while ensuring the safety of the citizens and preventing negative perceptions of large mammals.

While interactions with particular species result in adverse outcomes for people and/or biodiversity, the region also witnesses positive or neutral interactions between humans and biodiversity. Examples include the nesting of hornbills in human-dominated landscapes, the persistence of endangered tree species as individual trees in villages or within sacred groves (Blicharska et al 2013), and the preservation of aquatic ecosystems within sacred spaces (Aparna Watve pers. obs.). Such positive interactions need to be celebrated, documented, and revived. These can serve as examples for biodiversity and ecosystem conservation in other parts of the landscape.

#### Other threats

Konkan region of the Western Ghats receives over 3,000 mm of annual rainfall concentrated within the five-month monsoon period. Historically, this rainfall had supported semi-evergreen to evergreen forests (Biswas et al 2024). However, chronic anthropogenic disturbances, such as repeated clear felling and fires, have significantly altered plant composition. As a result, the landscape is dominated by species resilient to disturbance, such as *Terminalia elliptica* and *Terminalia paniculata*, which coppice easily and withstand fire. Fires are particularly widespread in the Ratnagiri District, where lopped tree vegetation is left to dry and then burned in fields, leading to uncontrolled fires that spread into adjacent forest patches (pers. obs. of multiple authors). These recurring fires, coupled with tree cover loss, have likely degraded soil properties (Verma and Jayakumar 2018), negatively impacting forest recovery. However, these impacts remain poorly studied and warrant urgent research attention and intervention. Additionally, the threats posed by rapid urbanisation and industrialisation, particularly downstream effects on local habitats due to industrial waste and pollution, in various towns in the region need to be thoroughly assessed to understand their ecological consequences. This may require conducting multi-taxa biodiversity assessments across the region to identify critical conservation areas that can guide future development activities.

### 4.3 Making information easily accessible

Interestingly, 77% of the literature we identified consisted of peer-reviewed publications or preprints, which have digital object identifiers (DOIs). We are reasonably confident that additional reports have been submitted to funding agencies or government departments. These reports likely contain vital information on biodiversity and conservation issues in the region, but are not readily available to citizens, environmentalists, funding agencies, or other scientists. Such information could help prevent duplication of effort, serve as evidence in court, and provide a baseline for future studies. These should be made publicly available on open-source platforms, such as Zenodo.

### 4.4 Role of non-governmental organisations in generating knowledge

While the role of environmental non-governmental organisations (NGOs) in environmental politics and discourse is recognised, their contribution to primary data generation is often under-appreciated (Partelow et al 2020; Rigolon and Gibson 2021). We found that NGOs play a significant role in generating knowledge about regional biodiversity. Given the common perception that NGOs primarily lead action-oriented projects, this finding underscores their crucial role in knowledge generation.

The under-representation of academic institutions in leading research efforts may be due to the Konkan region receiving less attention from academic institutions, as well as the likely perception that the northern Western Ghats are less ‘charismatic’ compared to the southern or central Western Ghats (as discussed during the focal group discussions). Unlike research institutions, colleges and universities often face heavier teaching loads, which constrain their ability to conduct ecological and conservation science research at sites far from their campuses. While academic institutions are mandated to conduct research, the primary focal areas of colleges, universities, and NGOs are teaching and/or conservation action. Such institutes could benefit from targeted capacity-building efforts in ecological methods and tools, further empowering them to conduct high-quality research. Higher education institutions have been advised to incorporate research as a key result area. However, its implementation has been weak.

Based on this study, we recommend the following: 1) conservation researchers can use the information in Table 2 to identify priority thematic areas for further research. These include topics of regional relevance with little existing information, such as linear intrusions, fires, and illegal wildlife trade, among others, 2) A horizon-scanning exercise to identify the most critical conservation questions for the region should be conducted to provide more specific guidance for future research, 3) Reports submitted to funding and government agencies should be made publicly available to enhance accessibility for relevant stakeholders and potentially indexed in a regional repository accessible to everyone, 4) conducting multi-taxa biodiversity assessments across the region to identify critical conservation areas which can be spared from anthropogenic disturbances, 5) identifying the primary socio-economic drivers of biodiversity loss in the region, and 6) considering the rapid growth of infrastructure, plantations, and tourism in Ratnagiri and Sindhudurg Districts, developing necessary policy guardrails to optimise biodiversity conservation and management in the region. The policy should be based on relevant findings from existing and future research, as well as consultations with regional stakeholders.

## ACKNOWLEDGEMENTS

We thank On the Edge Conservation, Godrej Consumer Products Limited and Gokhale Institute of Politics and Economics for their support. We thank Amaan Deshpande, Nikhil Atak, Pooja Sathe and Soomrit Chattopadhyay for their support during the workshop and Jithin Vijayan for his valuable inputs. We thank Anand Osuri, Kulbhushansingh Suryawanshi and Suri Venkatachalam for their helpful suggestions.

## Supplementary material

**Table S1:** List of keywords we used to search for literature on ecological and conservation science from the Konkan region. We used these keywords singly or in combinations to search for literature.

**Table S2:** Sources of information for rates of increase of certain human activities

**Table S3:** List of studies (published papers, preprints, reports) that we found during the literature survey.

